# A Frequency-Domain Framework for Cardiovascular Power Distribution

**DOI:** 10.1101/2025.07.31.667902

**Authors:** Abhishek Raje

## Abstract

Traditional cardiovascular models emphasize time-domain dynamics of blood flow and pressure, such as cardiac output and mean arterial pressure. However, the pulsatile nature of blood flow contains rich frequency content that interacts uniquely with organ-specific vascular properties. Drawing on wireless power transfer principles from electrical engineering, we propose a frequency-domain framework where the heart functions as a multi-frequency power source, generating a complex pressure waveform with multiple harmonics, and organs act as frequency-tuned loads, selectively absorbing power at their characteristic vascular resonance frequencies. This model introduces a frequency-division multiplexing analogy for cardiovascular power distribution, offering insights into physiological regulation, disease mechanisms, and therapeutic strategies. We present the theoretical foundation, mathematical models, physiological evidence, and potential clinical applications, supported by preliminary simulation results.

## I. Introduction

The cardiovascular system efficiently delivers oxygenated blood and nutrients from the heart to various organs, driven by pulsatile pressure waveforms. Traditional hemodynamic models focus on time-domain parameters like cardiac output and vascular resistance, often overlooking the frequency content of these waveforms. Each cardiac cycle produces a pressure waveform with a fundamental frequency (heart rate) and harmonics shaped by vascular reflections and waveform morphology.

Inspired by wireless power transfer (WPT) systems in electrical engineering, we propose a novel frequency-domain framework. Here, the heart is modeled as a multi-frequency power source, emitting a pulsatile waveform with multiple frequency components, while organs are conceptualized as frequency-tuned loads, absorbing energy at their vascular resonance frequencies. This analogy, akin to frequency-division multiplexing (FDM), provides a new perspective on cardiovascular power distribution, with implications for understanding physiology, diagnosing diseases, and developing therapies.

## II Multi-Frequency Wireless Power Transfer

### A. Overview of Wireless Power Transfer (WPT)

Wireless Power Transfer (WPT) transmits electrical energy without physical connections, often via magnetic resonance coupling. A transmitter coil generates an oscillating magnetic field, efficiently transferring power to a receiver coil tuned to the same resonant frequency, given by:

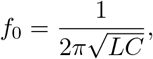

where *L* is inductance and *C* is capacitance. Resonance minimizes impedance, maximizing energy transfer efficiency [1].

### B. Frequency-Division Multiplexing (FDM)

In multi-frequency WPT, a single transmitter powers multiple receivers, each tuned to a distinct resonant frequency. This frequency-division multiplexing ensures selective energy delivery, minimizing interference. We extend this concept to the cardiovascular system, where the heart’s multi-frequency pressure waveform delivers energy to organs with unique vascular impedance profiles.

## III Novelty of the Frequency-Domain Approach

### A. Limitations of Current Models

Traditional cardiovascular models, such as lumped-parameter Windkessel models [2], focus on time-domain dynamics like pressure-flow relationships and cardiac output. While effective, these models underexplore the frequency content of pulsatile waveforms and their interactions with organ-specific vascular beds. Heart rate variability (HRV) analyses [4] assess autonomic function but do not model the heart as a dynamic, multi-frequency power source.

### B. Motivation for the Frequency-Domain Framework

Our framework models the heart as a variable-frequency power source, generating a pressure waveform with a fundamental frequency (heart rate, typically 1–2 Hz) and harmonics. Organs, with distinct vascular impedance spectra, act as frequency-tuned loads, absorbing energy at resonant frequencies. This approach:

- Integrates cardiac pulsatility and vascular resonance for a holistic view of power distribution.
- Accommodates HRV and waveform changes as modulators of spectral content.
- Offers insights into pathophysiological disruptions, such as altered vascular compliance.
- Leverages electrical engineering principles for interdisciplinary modeling.

## IV. Conceptual Framework

### A. Heart as a Multi-Frequency Power Source

The heart generates a pulsatile pressure waveform, decomposable via Fourier analysis into a fundamental frequency and harmonics. We adapt the McSharry et al. model [3] to simulate a synthetic electrocardiogram (ECG) driving cardiac dynamics:

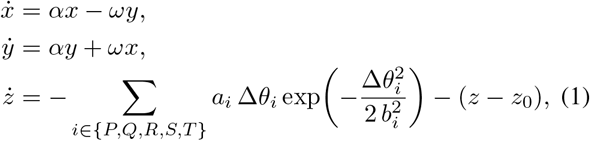

Where 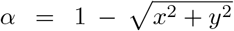, Δ*θ*_*i*_ = (*θ − θ*_*i*_) mod 2*π*, and *ω* is the angular frequency tied to heart rate. The *z*-dynamics generate a synthetic ECG (Fig. 1), reflecting the multi-frequency nature of cardiac output.

**Fig. 1:**
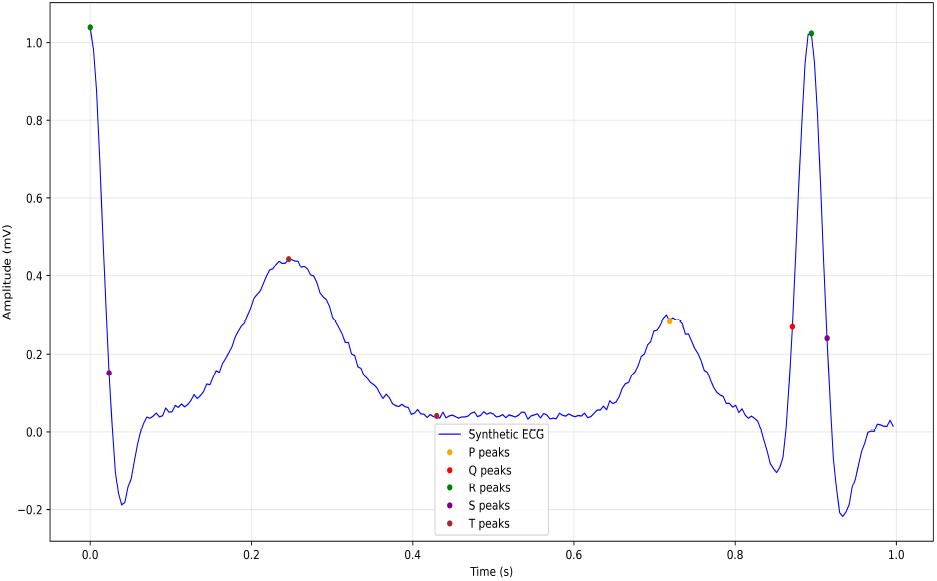
Synthetic ECG signal over one second, generated using the McSharry et al. model, illustrating the multi-frequency components of cardiac pulsatility.

To model HRV, we define the power spectrum of RR-interval variability:

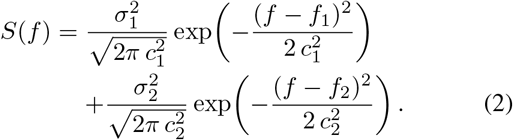

The RR-interval time series,*T* (*t*), is generated by taking the inverse Fourier transform of 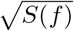 with random phases between 0 and 2*π* to capture frequency-domain variability. The parameter *ω* is varied for each oscillation according to this random process, as defined by the equation:

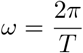

### B. Organs as Frequency-Tuned Loads

Organs are modeled as linear time-varying (LTV) RC circuits, with capacitance oscillating to mimic vascular compliance changes:

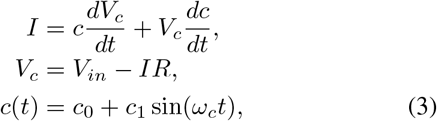

where *I* is current, *V*_*c*_ is capacitor voltage, *V*_*in*_ is input voltage, and *c*(*t*) is time-varying capacitance. The RMS power, calculated over the last 80% of the simulation, reflects energy absorption (Fig. 2).

**Fig. 2:**
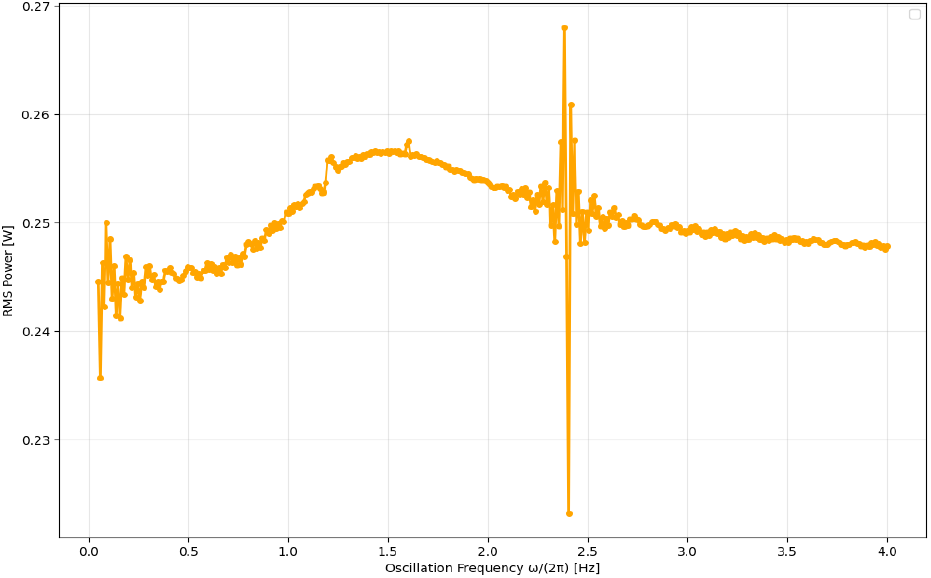
RMS power as a function of capacitance oscillation frequency for an RC circuit driven by a 1.2 Hz sinusoidal voltage, showing peaks at parametric resonance frequencies.

### C. Spectral Response and Parametric Resonance

Simulations using a 1.2 Hz sinusoidal input reveal parametric resonances at frequencies like 2*ω*, consistent with the Mathieu equation:

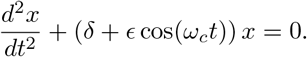

These resonances indicate efficient energy transfer at specific modulation frequencies. Fig. 3 shows RMS power as a function of capacitance oscillation frequency and simulation time, highlighting resonance-driven subharmonic energy peaks.

**Fig. 3:**
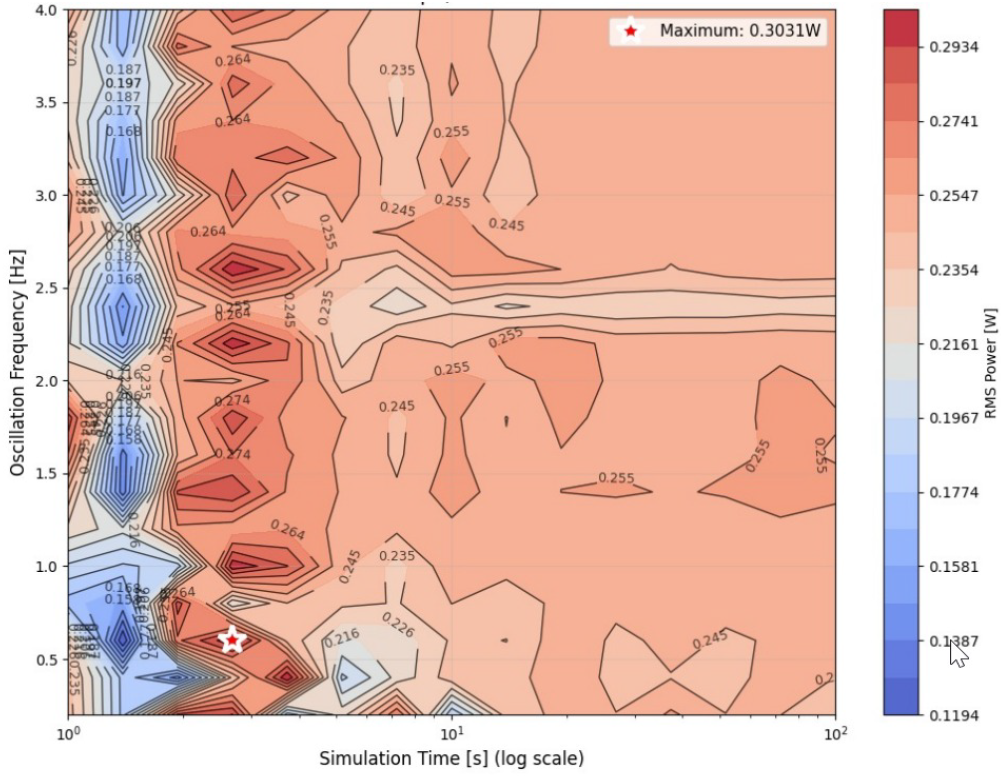
RMS power contour plot as a function of capacitance oscillation frequency and simulation time, illustrating frequency-dependent energy transfer.

### D. Energy Dynamics with ECG Modulation

We compared capacitor energy dynamics under sinusoidal and ECG-based capacitance modulation. The instantaneous energy is:

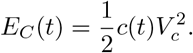

The RMS energy, calculated over the last 80% of the simulation, shows enhanced low-frequency activation under ECG modulation (Fig. 4), suggesting that the broadband nature of cardiac waveforms optimizes energy delivery to organs with low resonance frequencies.

**Fig. 4:**
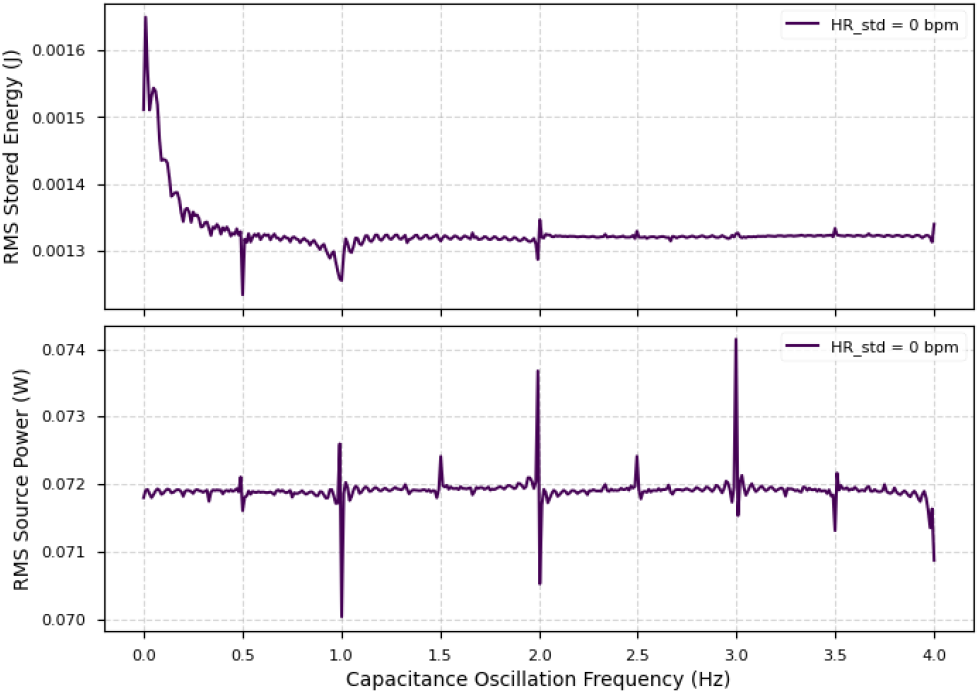
Top: RMS stored energy in the capacitor as a function of capacitance oscillation frequency under zero heart rate variability (HRV = 0 bpm). Bottom: Corresponding RMS source power. In the absence of HRV, the cardiac waveform is strictly periodic, leading to narrow spectral content. Energy absorption is reduced and constrained to discrete resonance points, demonstrating limited frequency engagement by the organ model

### E. Effect of HRV on Power Transfer

Heart rate variability (HRV) modulates the spectral content of the cardiac pressure waveform. As HRV increases, the beat-to-beat variability introduces a broader set of frequency components into the cardiac output.

This expansion creates multiple spectral bands across which energy is distributed. Organs with different resonance frequencies can absorb energy from the bands that align with their impedance characteristics.

Fig. 5 shows that as HRV increases from 5 to 15 bpm, both the RMS stored energy in the organ model and the RMS power drawn from the source increase across a range of capacitance oscillation frequencies. This confirms that higher HRV enhances power delivery by generating more frequency bands that organs can couple with.

**Fig. 5:**
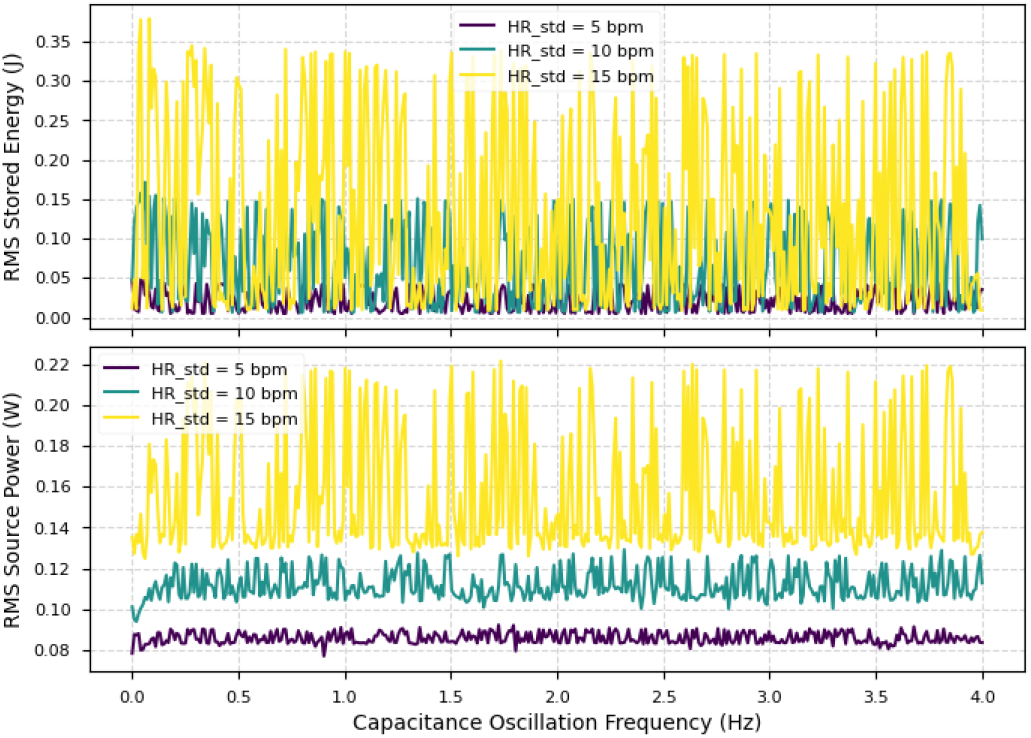
Top: RMS stored energy in the capacitor as a function of capacitance oscillation frequency for different HRV values. Bottom: Corresponding RMS source power. Increased HRV introduces more spectral bands, improving alignment with organ resonance frequencies and enhancing energy absorption.

This supports the interpretation of HRV as a spectral spreading mechanism that facilitates more effective power distribution to frequency-tuned organs.

## V. Breakdown of Resonant Relationship in Disease

### A. Vascular Stiffness and Resonance Shift

Cardiovascular diseases like hypertension and atherosclerosis increase arterial stiffness, reducing compliance (*C*) and shifting vascular resonance frequency:

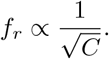

This reduces overlap with the heart’s frequency components, impairing power transfer and increasing cardiac workload [2].

### B. Altered Pulse Wave Dynamics

Disease-induced changes in arterial stiffness alter pulse wave velocity and reflection, distorting the harmonic content of pressure waveforms. This mismatch disrupts organ-specific energy absorption, contributing to microvascular dysfunction [5].

### C. Impaired HRV and Autonomic Dysfunction

Reduced HRV in diseases like heart failure limits the heart’s ability to modulate frequency content, disrupting resonance with vascular beds [4]. This impairs adaptive power distribution, exacerbating organ dysfunction.

### D. Supporting Evidence

Studies show increased arterial stiffness in hypertension [7] and disrupted cardio-renal coherence in chronic kidney disease [8], supporting the relevance of resonance disruption.

## VI. Future Directions

### A. Experimental Validation

Validate the framework by measuring organ-specific vascular impedance spectra using Doppler ultrasound and phase-contrast MRI. Correlate these with arterial pressure harmonics under varying conditions (e.g., exercise, hypertension) to confirm frequency-tuned power absorption.

### B. Computational Modeling

Develop models integrating time-domain hemodynamics with frequency-domain impedance spectra to simulate power distribution. Machine learning can enhance predictions of resonance mismatches in disease states [6].

### C. Clinical Applications

Use spectral analysis of arterial waveforms and HRV to detect resonance mismatches as biomarkers for early vascular dysfunction. Therapeutic strategies, such as pharmacological modulation of vascular tone or biofeedback to enhance HRV, could restore resonance and improve perfusion.

### D. Interdisciplinary Collaboration

Foster collaboration among physiologists, bioengineers, electrical engineers, and clinicians to refine models, develop diagnostic tools, and translate findings into clinical practice.

## VII. Conclusion

This frequency-domain framework redefines cardiovascular power distribution by modeling the heart as a multi-frequency power source and organs as frequency-tuned loads. Supported by simulations and physiological evidence, it offers a novel perspective on hemodynamic regulation and disease mechanisms. Future research and interdisciplinary efforts can translate this framework into diagnostic and therapeutic innovations, advancing personalized cardiovascular care.

